# A tensile ring drives tissue flows to shape the gastrulating amniote embryo

**DOI:** 10.1101/412767

**Authors:** Mehdi Saadaoui, Francis Corson, Didier Rocancourt, Julian Roussel, Jerome Gros

**Author notes:** To whom correspondence should be addressed, (JG); (FC). These authors contributed equally to this work (Mehdi Saadaoui, Francis Corson).

## Abstract

Tissue morphogenesis is driven by local cellular deformations, themselves powered by contractile actomyosin networks. While it is well demonstrated that cell-generated forces at the microscopic scale underlie a variety of local morphogenetic processes (e.g. lengthening/ narrowing^1–4^, bending^5–8^, or folding^9,10^), how such local forces are transmitted across tissues to shape them at a mesoscopic scale remains largely unknown. Here, by performing a quantitative analysis of gastrulation in entire avian embryos, we show that the formation of the primitive streak and the associated large-scale rotational tissue flows (i.e. ‘polonaise’ movements^11,12^) are integral parts of a global process that is captured by the laws of fluid mechanics. We identify a large-scale supracellular actomyosin ring (2 mm in diameter and 250 μm thick) that shapes the embryo by exerting a graded tension along the margin between the embryonic and extra-embryonic territories. Tissue-wide flows arise from the transmission of these localized forces across the embryonic disk and are quantitatively recapitulated by a fluid-mechanical model based on the Stokes equations for viscous flow. We further show that cell division, the main driver of cell rearrangements at this stage^13^, is required for fluid-like behavior and for the progress of gastrulation movements. Our results demonstrate the power of a hydrodynamic approach to tissue-wide morphogenetic processes^14–16^ and provide a simple, unified mechanical picture of amniote gastrulation. A tensile embryo margin, in addition to directing tissue motion, could act as an interface between mechanical and molecular cues, and play a central role in embryonic self-organization.

---

During amniote gastrulation, endodermal and mesodermal derivatives internalize through the primitive streak, a transient structure at the midline of the early embryo. In avians, the primitive streak forms from an initially crescent-shaped region at the margin between the embryo proper (EP) and extra-embryonic tissue (EE) (Fig. 1a), which converges towards and extends along the midline. While Myosin-II-driven oriented cell intercalation is known to underlie convergence-extension of the prospective primitive streak ^4,17^, how the concomitant vortex-like tissue flows arise and how they relate to the formation of the primitive streak has remained elusive, leaving our understanding of gastrulation incomplete.

**Figure 1.**
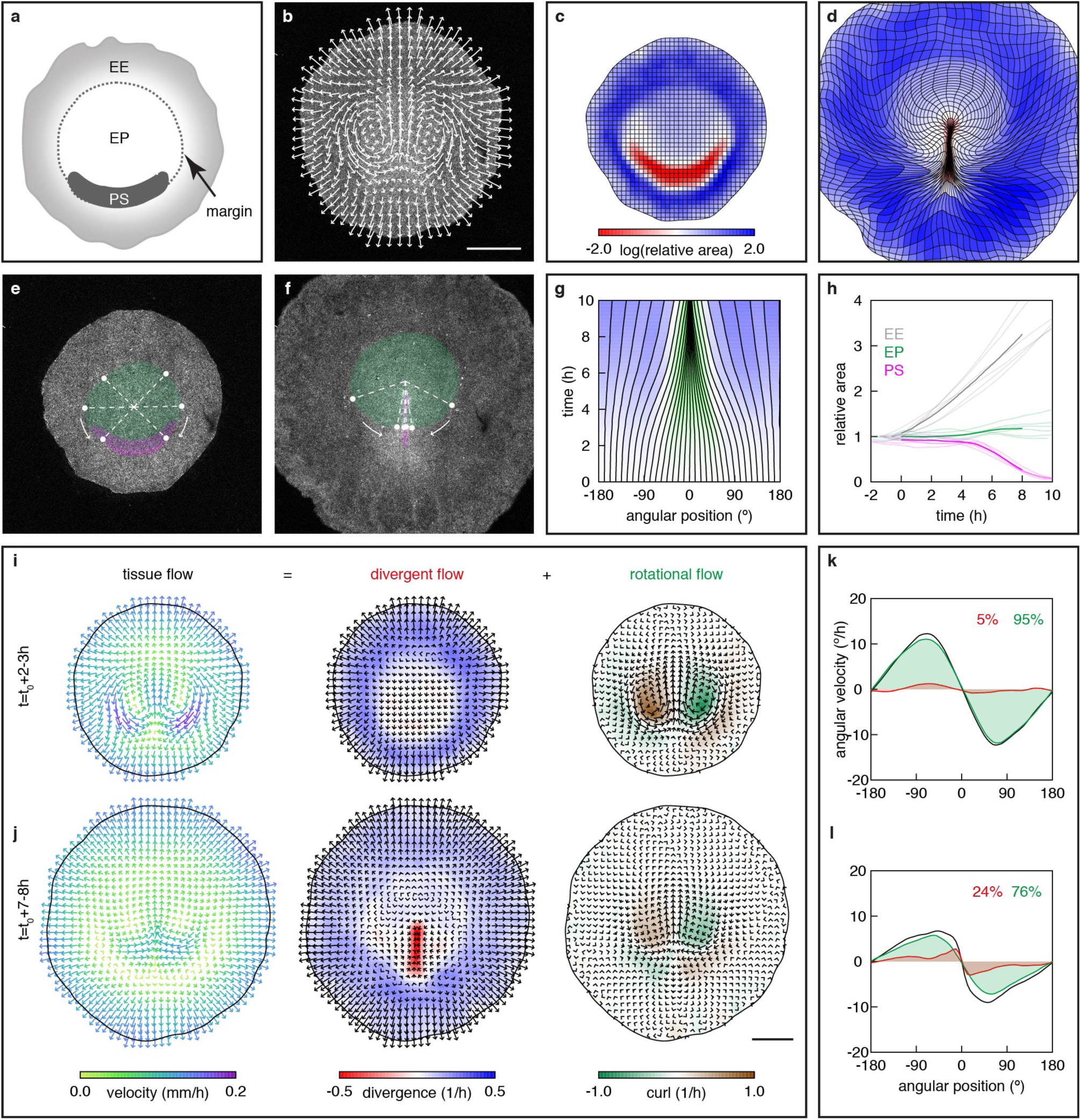
Quantitative description of gastrulation movements. **a**, Anatomical description of the early epiblast (EE, extra-embryonic territory; PS, prospective primitive streak; EP, embryo proper). **b**-**d**, Trajectories (**b**, *t* = 4-6 h) and deformation of an initially square grid (**c**, *t* = 0; **d**, *t* = 10 h), from the PIV analysis of a memGFP embryo movie (colors in **c**, **d** show area changes between the initial (**c)** and final (**d)** configurations; the primitive streak appears black in **d** due to denser grid lines). **e**-**h**, Automated fate mapping (green, EP; magenta, PS; **e**, *t* = 0 h; **f**, *t* = 10 h); dots show winding motion (arrows) along the margin, quantified in **g** by the time evolution of angular positions (dotted lines in **e**, **f**; 0° is posterior); **h**, area of tissue regions vs. time (*n* = 6 embryos; bold lines, averages). **i**-**l**, Decomposition of the tissue velocity field into divergent and rotational components (**i**, **j**), and contributions to motion along the margin (**k**, **l**; colors as in **i**; percentages quantify shaded areas) (averages over *n* = 6 embryos and the indicated time intervals). *t*0, time of motion onset. Scale bars, 1 mm.

To analyze gastrulation movements, we generated a transgenic quail line that ubiquitously expresses a membrane-bound GFP (memGFP)^18^, allowing the visualization of the ≃100,000 epithelial cells of the epiblast. Transgenic embryos were cultured *ex vivo*^19^ and imaged in their entirety for 12 hours. The resulting movies were processed using particle image velocimetry (PIV) to reconstruct cell trajectories and tissue deformation maps (Fig. 1b-d and Supplementary Video 1). Embryonic territories, originally characterized using anatomical or molecular criteria, could be recognized in these maps (compare Fig. 1a and 1c), and we designed automated fate mapping methods that identify and track these territories on the sole basis of tissue movement (Fig. 1e, f, see also Supplementary Methods and Extended Data Fig. 1). Based on these landmarks, we registered movies of 6 embryos in space and time to construct an average embryo (Supplementary Video 2, see also Supplementary Methods and Extended Data Fig. 1), used as a reference in the following. Importantly, the development of embryos in culture was virtually indistinguishable from an embryo imaged directly in the egg (Extended Data Fig. 2 and Supplementary Video 3).

This global analysis of tissue motion revealed that points all along the margin wind around the EP as they converge to the posterior (Fig. 1e, f) and their angular motion captured the progress of gastrulation (Fig. 1g). We further observed that the EP maintains an approximately constant area, while the EE tissue steadily expands (Fig. 1h), prompting us to distinguish area changes from other contributions to tissue movement. A decomposition into divergent (area changes) and rotational (incompressible) components indicated that gastrulation movements can be understood as the sum of three simpler flows: i) a radial, outward movement of the expanding EE tissue; ii) an area-preserving flow with two vortices within the embryo proper; and iii) at later stages, inward movement driven by areal contraction along the streak (Fig. 1i, j and Supplementary Video 4). Strikingly, while large-scale flows in the epiblast have been proposed to passively ensue from the deformation of the mesendoderm, propagating to the surrounding tissue^4,17,20^, we found that rotational movement persists after the mesendodermal crescent has converged onto the midline (Fig. 1i, j), and that areal contraction makes a limited contribution to continued movement towards the streak (Fig. 1k, l) - suggesting that other forces must be at play.

For a viscous fluid that is described by the Stokes equations, these forces could be derived from the Laplacian of the velocity field (Supplementary Methods). Applied to tissue flows in the epiblast, this suggested a pattern of tangential forces along the embryo margin, extending well into the anterior (Fig. 2a and Supplementary Video 4). In the case where flow is driven by active internal stresses - here, by cell contractility -, forces inferred in this way should be understood as apparent external forces, arising from the spatial variations (the tensor divergence) of the active stress. The force pattern of Fig. 2a thus pointed to a tensile margin around the EP, with a tension that decays from posterior to anterior; tissue along the sides of the EP is drawn towards the posterior, where tension is higher; tissue in the posterior is thrust forward, because the margin is curved (Fig. 2b). To test whether such a tensile margin and a fluid-like tissue behavior could account for gastrulation movements, we formulated a mathematical model that is based on the Stokes equations for viscous flow. Motion in the model is the resultant of area changes, treated as the non-uniform growth of an otherwise incompressible tissue, and active tensile stresses along the margin, which moves with the tissue (Fig. 2c, d). Area changes are taken from experiment (individual or average embryo), while tensions along the margin are fit to the observed motion at each time step (Fig. 2c). Although strongly constrained -aside from the tensions, the initial position of the margin and its width are the only free parameters -, the model recapitulates the full course of tissue movements over 8 h (Supplementary Video 5), with 85-90% accuracy for individual embryos, and > 90% for the average reference embryo (normalized step-by-step residuals; Fig. 2f, g and Methods). The model can predict the full deformations that would result from each source term taken separately (Fig. 2e), and argues that active tensions largely account for the shaping of the embryo, while area changes are mostly responsible for EE expansion.

**Figure 2.**
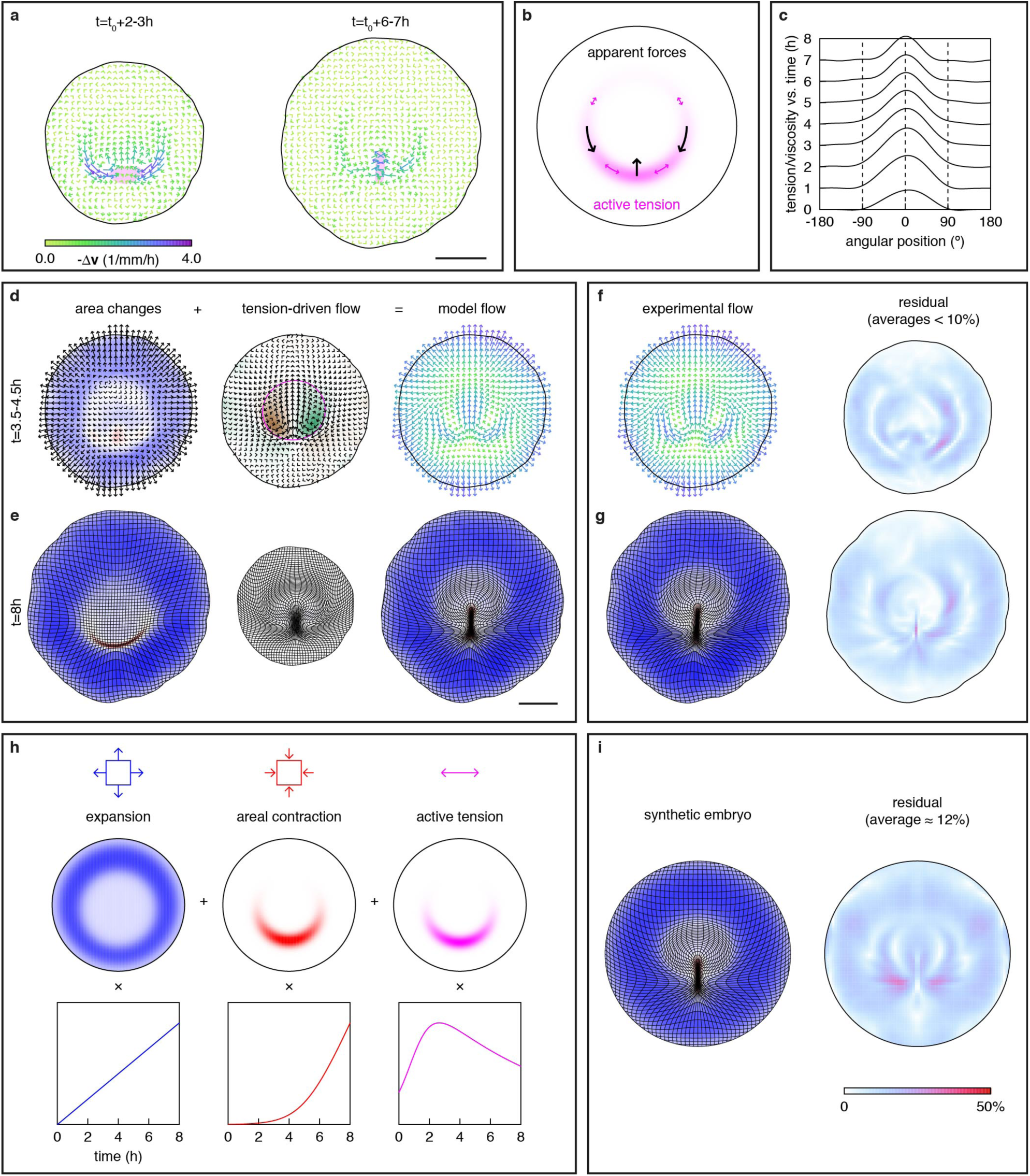
A quantitative fluid-mechanical model for gastrulation. **a**, Apparent forces (negative of the Laplacian of the velocity field; averages over *n* = 6 embryos and the indicated time intervals; magenta, presumptive primitive streak). **b**, Sketch illustrating how apparent forces (black) arise from graded tensions along the margin (magenta). **c**-**e,** Quantitative model for gastrulation movements. **c**, Tension/viscosity profiles (mm/h; averages over 1 h intervals) from a fit to the reference average embryo (*n* = 6 embryos). **d**, **e**, Tissue flows in the model as the resultant of area changes (taken from experiment) plus an incompressible flow driven by tension along the margin (magenta line in **d**). **d**, Velocity fields (colors as in Fig. 1i). **e**, Deformation maps from each source term taken separately and together. **f-g,** Velocity field (**f**) and deformation map (**g)** for the average embryo (right-hand panels show deviation between model and experiment). **h**, EE expansion, areal contraction of the prospective primitive streak, and tension along the margin as a function of space and time used to build a synthetic model of gastrulation. **(i)** Deformation map for the synthetic model and deviation from average embryo. Scale bars, 1 mm.

As a further abstraction, we constructed a “synthetic embryo”, where the three driving terms, EE expansion, areal contraction along the streak, and graded tension along the margin, are described by simple mathematical functions of space and time (Fig. 2h, Supplementary Video 6, see also Supplementary Methods, Extended Data Table 1 and Extended Data Fig. 3). Although reduced to minimal ingredients, this synthetic model captures the movement of the average embryo from which it was derived with 88% accuracy (normalized end-to-end residual; Fig. 2i and Methods). The model also lends itself to an analytical treatment, which recovers the essential features of the movement in the limit of a thin margin (Extended Data Fig. 4 and Supplementary Methods).

Our results so far demonstrated that gastrulation is best understood as a tissue-wide process and pointed to (i) a tensile margin and (ii) a fluid-like response of the epithelial epiblast as its major components. Further experiments were designed to substantiate these two components, and to challenge a fluid-mechanical description of gastrulation movements. First, to directly probe the existence of a tensile margin, we performed circular UV-laser cuts^21^ at different locations in the epiblast (Fig. 3a-c and Supplementary Video 7). Cuts in the anterior and posterior margin revealed anisotropic tissue strains, as expected. Cuts inside the EP showed significantly lower strains, which can be interpreted as a manifestation of passive viscosity (see Supplementary Methods). To connect tissue-scale motion and cellular-scale behaviors, we analyzed fixed embryos that had previously been live-imaged for different time intervals (Fig. 3d-m and Extended Data Figs. 5-8). Cell segmentation of entire embryos showed a gradual increase in cell areas in the EE tissue, likely contributing to its expansion (Fig. 3e, h, j, m). Cell shapes, with an initially isotropic distribution throughout the tissue, became elongated along the margin, consistent with a state of tension (Fig. 3f, h, k, m). Quantification of junctional phosphorylated Myosin II revealed localized myosin anisotropy, a correlate of active force generation^14^, at the margin (Fig. 3g, h, l, m). To obtain a more dynamic picture, we generated a transgenic quail line expressing a ubiquitous LifeAct-NeonGreen-ires-tdTomato-MyosinII transgene reporting on Actin and Myosin II dynamics. High-resolution live imaging revealed the progressive formation of dynamic, tangential actomyosin supracellular cables, spanning 5-20 cells specifically at the margin (Fig. 3n and Supplementary Video 8). In the posterior margin, these cables contracted, driving oriented intercalations^4^. In the anterior margin, supracellular cables were also visible but extended tangentially, concomitant with cell elongation and oriented divisions (Fig. 3o, p and Supplementary Video 9), indicative of stress dissipation^22^. Altogether, these results demonstrate that the margin exerts active tension in the posterior, and passive tension - possibly through locally enhanced resistance to stretching - in the anterior.

**Figure 3.**
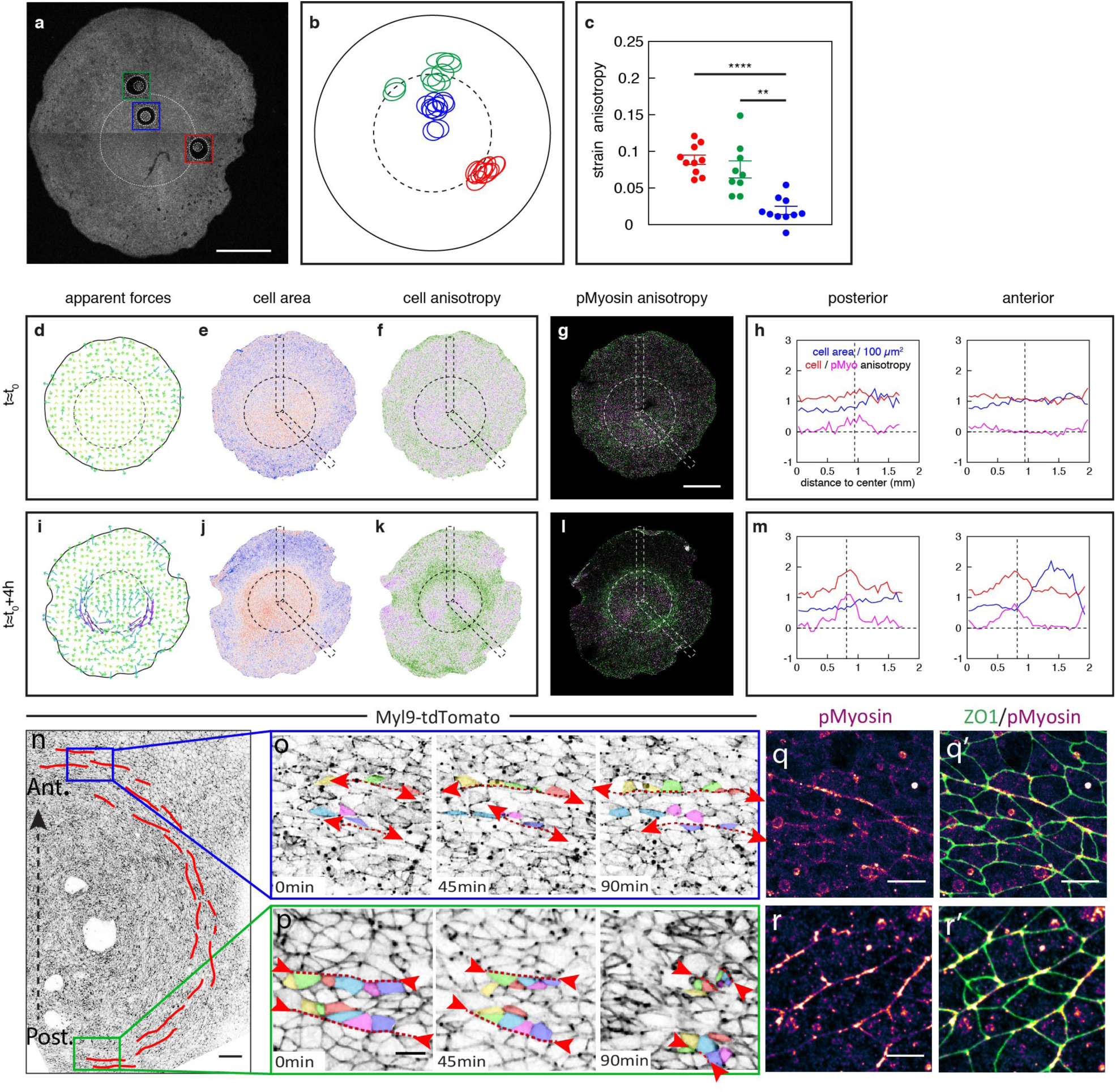
Mechanical, cellular, and molecular characterization of the embryo margin. **a**-**c**, 250 μm circular laser cuts in a single memGFP embryo (**a**; overlays show response at 2 min) and representation of all laser cut experiments (**b**; ellipses show cut location relative to margin and anisotropic strain, amplified 4-fold for visibility) for which the response was quantified (**c**; tangential vs. radial strain anisotropy based on stretch ratios along the two axes; bars, mean ± SE; ** P<.01; **** P<.0001; paired t-test) (red, posterior margin; green, anterior margin; blue, EP). **d**-**m**, Apparent forces (**d**, **i**) in embryos imaged until indicated stages (dashed line, margin), and cell areas (**e**, **j**), cell shape anisotropy (**f**, **k**; tangential vs. radial elongation), and junctional phosphorylated Myosin anisotropy (**g**, **l**) mapped for all cells in these embryos and quantified for two anterior and posterior radial boxes (**h**, **m**). **n**-**p**, Time series from Supplementary Video 8 and 9 showing supracellular actomyosin cables (red) at the margin in a Myl9-tdTomato transgenic embryo and concomitant cell behavior (tracked colored cells) at the anterior (**o**) and posterior (**p**) margin. Actomyosin cables extend anteriorly and contract posteriorly (arrowheads). **q**-**r**, Antibody staining for phosphorylated Myosin and ZO-1 in the anterior (**q**, **q’**) and posterior (**r**, **r’**) margin revealing orthoradial supracellular cables. Scale bars, **a**, **g**, 1 mm; **n**, 100 µm; **o**, **p**, 20 µm; **q**-**r**,10 µm.

Second, we sought to identify the cellular basis of tissue fluidity. Since cell rearrangements contribute to stress relaxation in epithelial tissues^23^, and since most cell rearrangements in the early avian embryo are associated with cell division^13^, we reasoned that cell division may be required for a fluid-like behavior. Treatment with hydroxyurea (HU) efficiently suppressed cell division (Fig. 4a), but, like every other cell division inhibitors we tested (not shown), it induced apoptosis on the long term (Extended Data Fig. 9a). To circumvent this toxicity, HU was combined with Q-VD-OPh, a potent apoptosis inhibitor. In these conditions, both cell division and apoptosis were suppressed (Fig. 4a and Extended Data Fig. 9b, c), and the topology of the epithelium was greatly stabilized compared to control embryos (Fig. 4b, c and Supplementary Video 10). While Q-VD-OPh alone had no noticeable effect on cell dispersal, and only slightly delayed the progress of gastrulation movements (Extended Data Figs. 1j and 9d-h), embryos incubated in both HU and Q-VD-OPh showed a dramatic slowdown by 6-8h of treatment and failed to form a primitive streak (*n* = 6, Fig. 4d, e, g, Extended Data Fig. 1j and Supplementary Video 11). Strikingly, whereas tissue expansion persisted, rotational movements were abolished, consistent with a suppression of fluidity (Fig. 4f and Supplementary Video 12). Based on the model, which fit treated embryos equally well as control embryos, the amplitude of the tension/viscosity ratio dropped over time (Fig. 4h). Laser cuts in embryos treated with HU and Q-VD-OPh revealed that tensions were still present, if not increased, along the margin (Extended Data Fig. 9i-k and Supplementary Video 13) - implying that the slowdown resulted from an increase in viscosity and not from a decrease in tension. Together, these results demonstrate that the fluidity of the embryonic epithelium is required for primitive streak formation and indicate that it emerges from cell division.

**Figure 4.**
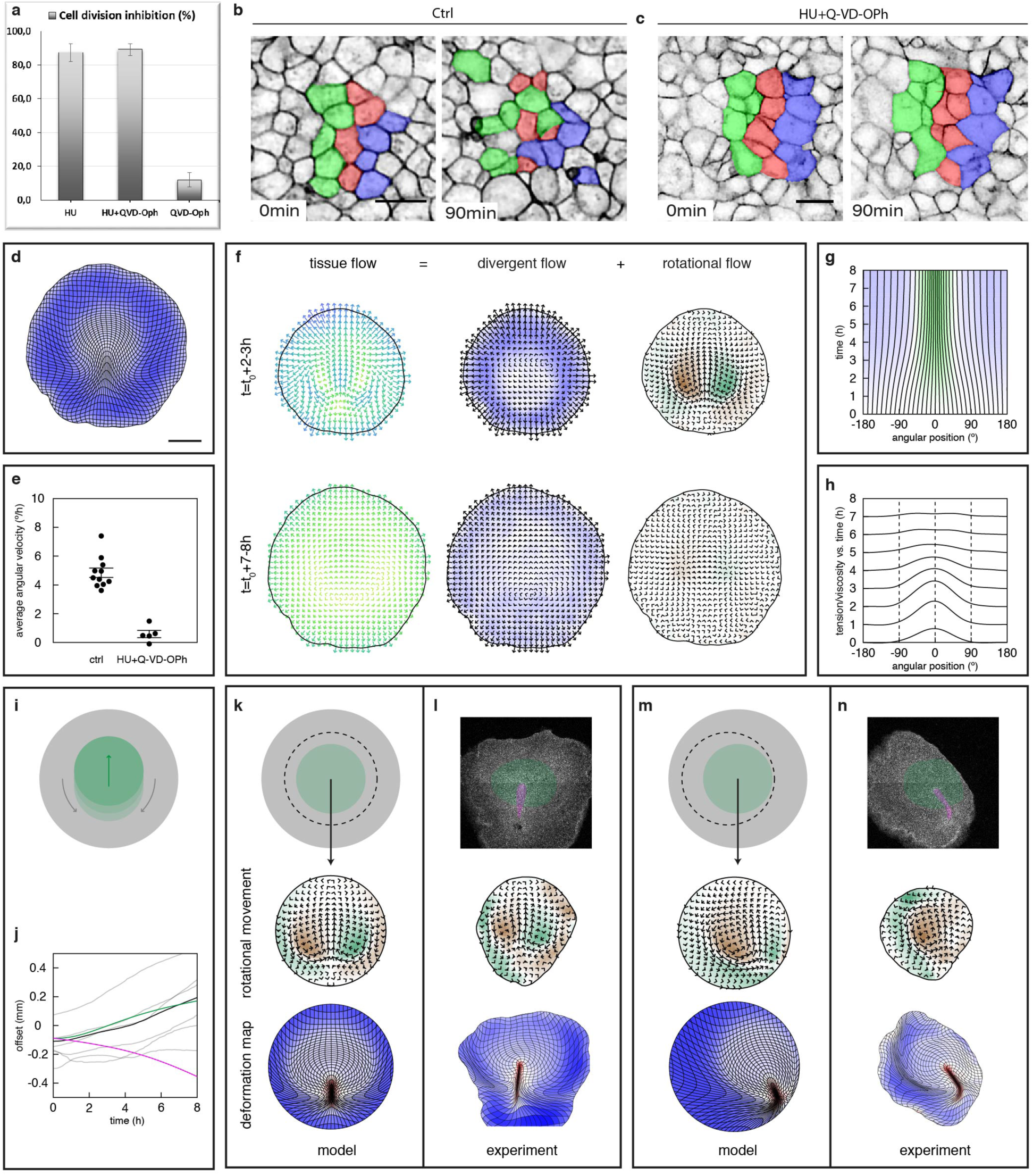
Manipulation of tissue viscosity and hydrodynamic effects in gastrulation. **a**, Percentage of cell division inhibition in drug treated embryos. **b**, **c,** Effect of cell division on cell rearrangements revealed by cell dispersion in control (**b**) and treated (**c**) embryos. **d**-**h**, Effect of HU+Q-VD-OPh on gastrulation movements (*n* = 5 embryos). **d**, Deformation map (average embryo). **e**, Average angular velocity along the margin at *t* = *t*_0_ +7 h. **f**, Decomposition of the tissue velocity field into divergent and rotational components (averages over embryos). **g**, Time evolution of angular positions along the margin (average embryo). **h**, Tension/viscosity profiles from model fit to average embryo. **i**, Sketch of swimming motion (green arrow) induced as the EP (green) draws EE tissue (gray) posteriorly (gray arrows). **j**, Anterior movement of embryos relative to tissue border (gray, *n* = 6 embryos; black, average embryo; green, synthetic embryo; magenta, synthetic embryo without active tensions). **k**-**n**, Predictions from the synthetic model (**k**, **m**) and experimental response to centered (**k**, **l**) and off-centered (**m**, **n**) cuts generating a new tissue border. Scale bars, **b**, **c**, 20 µm; **d**, **l**, 1 mm.

Third and finally, we challenged model predictions for hydrodynamic effects in gastrulation. The embryo, which draws the surrounding tissue to the posterior, is akin to a swimmer, and should move forward over time (Fig. 4i). Indeed, embryos exhibited a slow anterior-ward movement, in quantitative agreement with the model (Fig. 4j); in the absence of active tensions, the initially off-centered embryo is predicted to drift posteriorly, i.e. the embryo swims against a current associated with tissue expansion. At odds with the view that vortex-like flows are shaped by a confining boundary^20,24,25^, our model suggests that they are governed by the distribution of active forces, with boundary conditions playing a limited role. The progress of gastrulation is predicted to be weakly sensitive to the distance to epiblast border, reaching > 80% of its maximum rate when it is just 50% larger in radius than the EP (Extended Data Fig. 4d and Supplementary Methods). Indeed, circular cuts centered on the margin, that removed most of the EE tissue and brought the epiblast border closer to the EP, had almost no effect on tissue flow and streak formation (Fig. 4k, l and Supplementary Video 14). By contrast, in the case of off-centered cuts, which bring the border even closer to one side of the EP, the model predicted, and experiments confirmed, that the interaction between EP and border induces a rotation of the axis, leaving only one apparent vortex, and resulting in a bent streak (Fig. 4m, n and Supplementary Video 13). While the predictions are inevitably qualitative, the model fit *a posteriori* to these strongly perturbed embryos recapitulated their motion, with an accuracy that is comparable to controls. These experimental validations demonstrate the predictive power of our model and the relevance of a fluid-mechanical description of gastrulation.

Our global mechanical analysis of gastrulation demonstrates that tissue-wide flows in the embryonic disk are not simply a passive consequence of primitive streak formation. Instead, both are part of a broader process, driven by a steady pattern of forces around the margin, which shapes the embryo as a whole. If mechanical forces feed back onto themselves and onto gene expression, a tensile embryo margin could play a central organizing role in the establishment of the amniote body plan.

## Methods

### Animals

All experimental methods and animal husbandry procedures to generate transgenic quails were carried out in accordance with the guidelines of the European Union 2010/63/UE.

### Production and characterization of transgenic quail lines

Two transgenic lines were created in this study (*hUbC:memGFP* and *hUbC:Lifeact-NeonGreen-ires-Myl9-tdtomato)* by following a previously published method ^26^. Briefly, non-incubated quail eggs (*Coturnix japonica*) were windowed and a solution of high titer lentivirus was injected into the subgerminal cavity of stage X embryos. Eggs were sealed with a plastic piece and paraffin wax. Injected eggs were incubated at 37.5°, 56% humidity until hatching. For the *hUbC:memGFP* line, a total of 42 embryos were injected with the lentivirus solution (titer 10^10^/ml). Three F0 mosaic founder males successfully hatched and reached sexual maturity (7%). They were bred to WT females and all three produced transgenic offspring (transmission rate: 8.8%). One line was selected on the basis of a single copy of the transgene, checked by Southern Blot, and high intensity of the memGFP signal. For the *hUbC:Lifeact-NeonGreen-ires-Myl9-Tdtomato,* a total of 92 embryos were injected with the lentivirus solution (titer 4.5 10^7^/ml). Four F0 mosaic founder males hatched, reached sexual maturity (4%) and were bred to WT females. One line with a single copy of the transgene was segregated, by Southern Blot analysis, from a line presenting three copies of the transgene and a high intensity of the tdTomato signal.

### Embryo culture, time-lapse microscopy and laser severing experiments

Transgenic quail eggs were collected at stage XI and cultured using a modified version of the EC culture system ^16^ until stage 4. Briefly, embryos were collected using paper filter rings and cultured on mix of albumen, agarose (0.2%), glucose, and NaCl. Embryos were then transferred into a bottom glass Petri dish (Mattek inc.) with semi solid albumen/agarose nutritive substrate with or without drugs: HydroxyUrea (40mM), Q-VD-OPh (250µM-1mM) for imaging. Embryos were then imaged at 38°C using an inverted confocal microscope (Zeiss LSM 880 or LSM 700) or a 2-photon Microscope (Zeiss, NLO LSM 7MP) coupled to a Chameleon Ti/Saph femtosecond pulsed laser (Coherent inc.) at 840nm wavelength using 5X, 10X or 40X long distance objectives.

Laser microdissections were performed during image acquisition using a 355-nm pulsed laser (75-100% power), a UGA-42 module from Rapp Optoelectronic coupled to a Zeiss LSM 880 and a 5X or 10X objective. Before and after severing, the time interval between two consecutive frames was 6 min. Briefly, a PIV analysis as described below was used to define ROIs of different sizes in order to isolate the embryonic region (from 1.8mm to 2.5mm) or to measure the tension inside the epiblast (between 250 and 500µm). Images were acquired during severing every 5s. The tissue strain was evaluated based on the deformation of the tissue 2 min after the cut, from a PIV analysis of the resulting time-lapse movies (see Supplementary Methods for details; relaxation was sufficiently fast, compared to the severing time, that the initial relaxation rate could not be measured).

### Quantitative analysis of tissue flows

Time-lapse movies of embryos were analyzed using custom Java software, building on the ImageJ API for image processing. The full details of processing and analysis are given in Supplementary Methods. Briefly, particle image velocimetry (PIV) was used to evaluate the local displacement of the tissue between successive movie frames. The resulting displacement fields were used to reconstruct cell trajectories, and a family of mappings that relate the configuration of the tissue at any two time points. In terms of these mappings, deformation maps as in Fig. 1d show the image of an initially square grid at a later time. Tissue velocity fields were decomposed into a divergent, approximately irrotational component, and a divergence-free, rotational component using a variation on the Helmoltz-Hodge decomposition of vector fields, adapted such that the divergent component coincides with the full velocity field at the tissue border (this implies that, in general, the divergent component is not exactly irrotational).

### Automated fate mapping and spatiotemporal registration

The location of the embryo margin was identified using an active contour or “snake” approach^27^; as a first step, we constructed a function of space that changes sign across the margin, based on the distinctive tissue deformation patterns observed on either side. The primitive streak was identified from the pattern of areal contraction in the tissue, by thresholding. The orientation of the posterior was identified as a fixed point in the angular motion of points along the margin. Embryos were registered in space based on the location of the margin, and in time based on the time of motion onset and the time of streak formation, both defined according to the angular contraction of an initially 90° sector centered on the posterior (see Supplementary Methods for details).

### Fluid-mechanical model

The full details of model derivation, simulation, fitting, and analysis, are given in Supplementary Methods. Briefly, the tissue is treated as effectively incompressible, and areal expansion or contraction as intrinsic behaviors of different tissue regions, akin to growth. Taking the long-time limit of a viscoelastic description of the tissue, we recover the Stokes equations for viscous flow, with an additional source term for areal expansion or contraction,

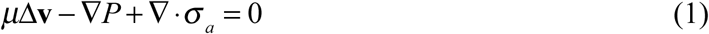

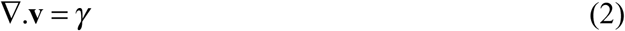

Equation (1), with **v** denoting the velocity, *μ* the viscosity, *P* the pressure, and *σ* _a_ an active internal stress tensor, expresses force balance, and is unchanged from the incompressible fluid case. In equation (2), the usual incompressibility condition ∇ · **v** = 0 is replaced by a prescribed divergence *γ*, corresponding to the local rate of areal expansion or contraction. Area changes and the motion of the tissue border, which defines the boundary condition of the model, are taken from experiment (from an individual or average embryo). The active internal stress takes the form of a tensile stress along the margin, represented by a contour that is advected with the tissue. The tensile stress has a Gaussian profile across the width of the margin, and its magnitude at different positions along the margin is represented by the total tension across the width of the margin. For an approximately circular margin, the average tension has almost no incidence on motion, and it is arbitrarily chosen such that the tension vanishes in the anterior. Tension profiles at each time step are fit to the observed tissue motion, leaving the initial position of the margin and its width as fitting parameters. The source term for area changes, *γ*, is computed from the observed tissue flow in two different ways. In the simplest, “Eulerian” treatment, it identifies with the divergence of the observed velocity field, and the divergent component of the model flow coincides with its experimental counterpart. In a refined, “Lagrangian” treatment, the rate of areal expansion or contraction is a property of material points in the tissue, and thus tied to the initial, rather than the current configuration of the tissue. The “Eulerian” treatment is used for parameter fitting. The “Lagrangian” model, which is computationally more expensive, is simulated with the resulting parameter values, as the definitive form of the model. The synthetic model is obtained by replacing the source terms - area changes and tensions - by analytical functions of space and time. As a boundary condition, the tissue border is circular and moves outwards at a uniform velocity, determined at each time step according to the integrated area change. The resulting model, which is left-right symmetric by construction, was fit and compared to a symmetrized average embryo, obtained by including a mirror image of each embryo.

### Model residual

Two measures of the deviation between model and experiment are reported in the text. A normalized step-by-step residual is obtained as the magnitude (the *L*^2^ norm) of the deviation between the displacement fields at each time step in model and experiment, *δ* **r** and *δ* **r**_exp_, divided by the magnitude of the experimental displacements,

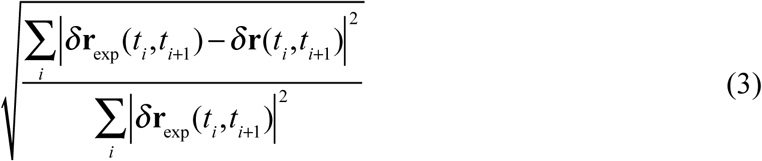

A normalized end-to-end residual is defined in the same way from the displacements between the initial and final time points,

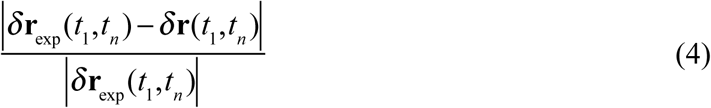

Step-by-step residuals for *n* = 6 control embryos, evaluated using 1 h steps to limit the contribution of fluctuations, ranged from 10.4 to 15.5%.

### Analysis of cell behaviors and fixed embryos

Time-lapse movies were analyzed using Fiji^28^ and Icy software. Quantifications were performed manually on registered movies ^29^ by visual inspection. For cell division quantification, ROIs from at least 3 independent experiments were used to count the number of dividing cells in the first 5-6h of the movies. For epithelial stability in different conditions (ctrl, HU, HU+Q-VD-OPh, and Q-VD-OPh) and after a step of image segmentation (described below), the Epitools^30^ plugin for Icy^31^ was used to generate the movies with the cell color tag option in the cell overlay panel.

For fixed samples, optical sections were obtained on a confocal microscope (LSM700 or LSM880; Zeiss) using a 40X (C-Apochromat NA 1.1 water immersion) objectives and Zen software (Zeiss). Images were analyzed using Fiji and Icy softwares. For segmentation, the apical signal was first extracted from image stacks using a custom program that fits a smooth surface to the tissue. After image processing, a binary mask was generated using the Find Maxima tool of Fiji and then subjected to manual correction using Tissue Analyzer^32^. The segmented images were then analyzed as described in Supplementary Methods.

### Immunofluorescence

For antibody stainings, quail embryos were fixed in ice cold 4% formaldehyde/PBS for at least 1h, permeabilized in PBS/0.1% Triton X-100 (PBT 0.1%) before a blocking step in PBT 0.1%/2% BSA (from Roche)/10% FBS (from Gibco). Primary antibodies used in this study are mouse anti-ZO1 (Invitrogen ZO1-1A12), rabbit anti-pMyosin light chain 2 (Cell Signaling Technology CST-3671S and CST-3674S), mouse anti-β-Catenin (BD Transduction Laboratories™, clone 14) and rabbit anti-h/mCaspase3 (RD Systems AF835). Secondary antibodies coupled to AlexaFluor 488, 555, or 647 were obtained from Invitrogen and used at 1:200 dilutions. Embryos were then mounted with DAPI-containing Fluoromount-G^™^ (eBioscience) between slide and coverslip.

## Acknowledgements

We thank Vincent Hakim, Pierre-Francois Lenne, Alfonso Martinez Arias, and François Schweisguth for critical reading of the manuscript. The research leading to these results has received funding from the European Research Council under the European Union’s Seventh Framework Programme (FP7/2007-2013) / ERC Grant Agreement *n°337635,* from the Institut Pasteur, the CNRS, the Cercle FSER, the Fondation pour la Recherche Medicale.

## Author Contributions

M.S., F.C. and J.G. conceived the study. D.R. and J.R. generated transgenic lines. M.S. performed experiments. F.C. developed quantitative analysis methods and theoretical models. F.C. and J.G. wrote the paper with input from M.S.

## Extended Data

**Extended Data Figure 1.**
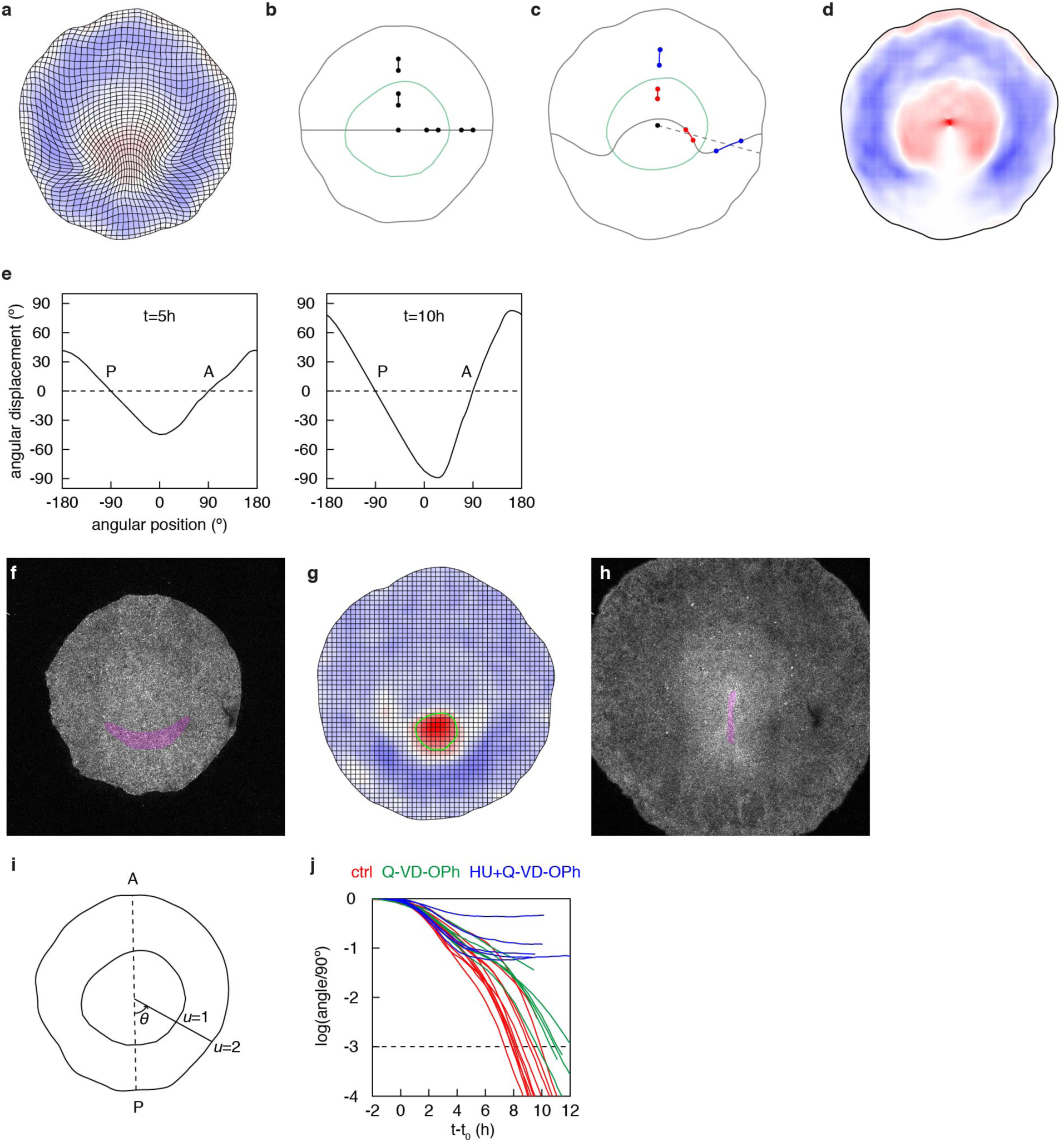
Automated fate mapping and spatiotemporal registration. **a**-**d**, Embryo proper. **a**, Deformation of the tissue over a 4 h interval. **b, c**, An initially straight line, and segments on either side of the margin (green) are tracked over the same time interval, showing: on the right-hand side, differential rotation relative to a radial line (dotted line in **c**); in the anterior, differential length changes (red and blue denote interior and exterior as in **d**). **d**, Plot of the criterion function used to identify the embryo margin (red, interior; blue, exterior). **e**, Anterior-posterior axis. Plots of the angular displacement of points along the embryo margin, as a function of initial angular position (relative to the horizontal image axis), over intervals of 5 and 10 h. The anterior (A) and posterior (P) are identified as zero crossings of the angular displacement. **f**-**h**, Prospective primitive streak. **f**, **h** are two images of an embryo at the onset of motion and at the “streak stage”, respectively, while **g** shows the area changes between an intermediate stage and **h**, similar to Fig. 1c. The boundary of the prospective primitive streak is identified by thresholding the area changes (green line in **g**). The corresponding regions are highlighted in **f**, **h** (magenta). **i**, Coordinate system for spatial registration. **j**, Contraction of a 90° posterior sector vs. time, used for temporal registration, in control and treated embryos (the dashed line defines the “streak stage”).

**Extended Data Figure 2.**
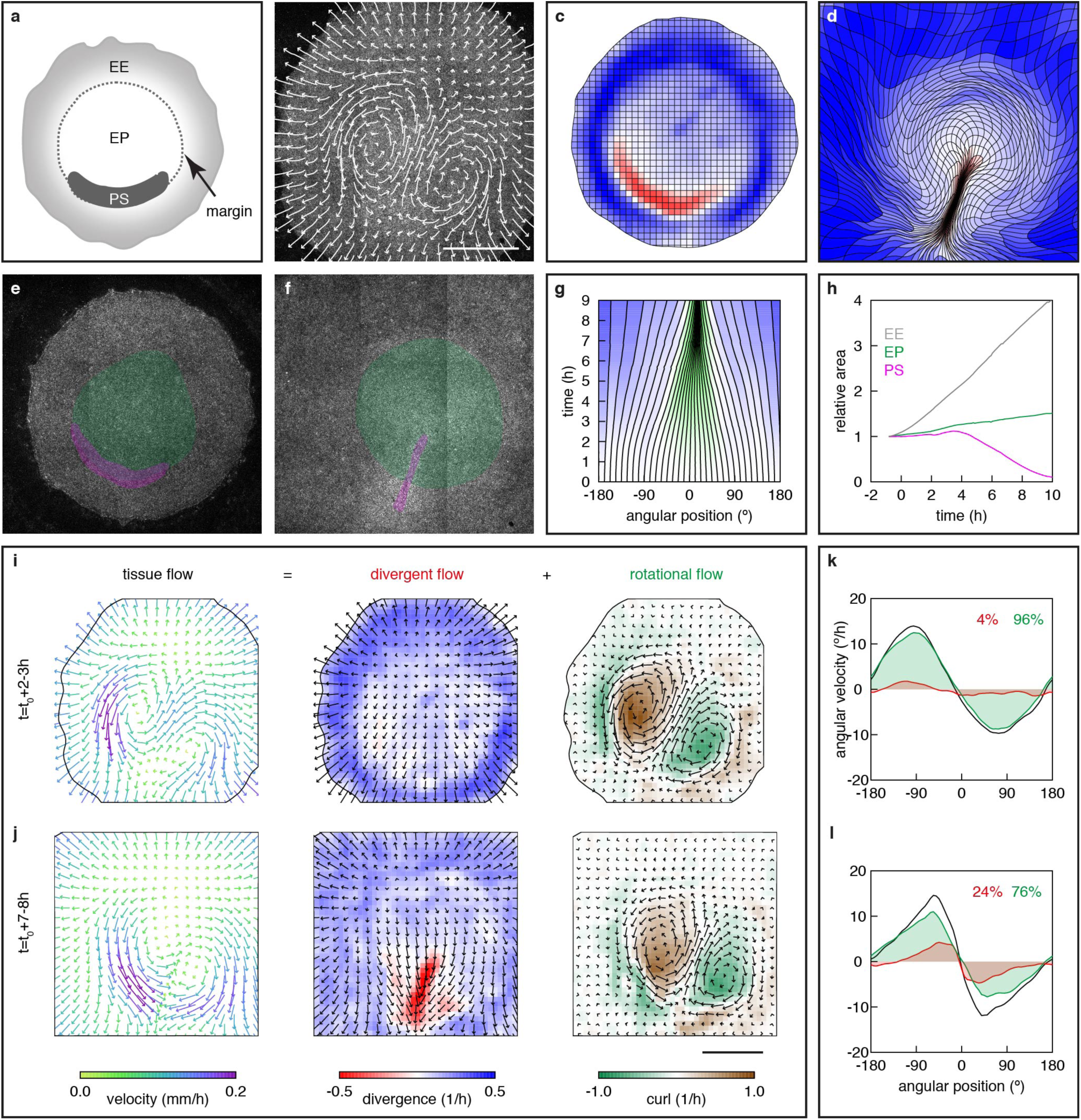
Quantitative description of gastrulation movements of an embryo imaged directly in the egg, without *ex vivo* culture. (Similar analysis as in Fig. 1). **a**, Anatomical description of the early epiblast (EE, extra-embryonic territory; PS, prospective primitive streak; EP, embryo proper). **b**-**d**, Trajectories (**b**, *t* = 3.5-5.5 h) and deformation of an initially square grid (**c**, *t* = 0; **d**, *t* = 9 h), from the PIV analysis of a memGFP embryo movie (colors in **c**, **d** show area changes between the initial (**c)** and final (**d)** configurations). **e**-**h**, Automated fate mapping (green, EP; magenta, PS; **e**, *t* = 0 h; **f**, *t* = 9 h); **g,** time evolution of angular positions; **h**, area of tissue regions vs. normalized time. **i**-**l**, Decomposition of the tissue velocity field into divergent and rotational components (**i**, **j**), and contributions to motion along the margin (**k**, **l**) (averages over the indicated time intervals). Scale bars, 1 mm.

**Extended Data Figure 3.**
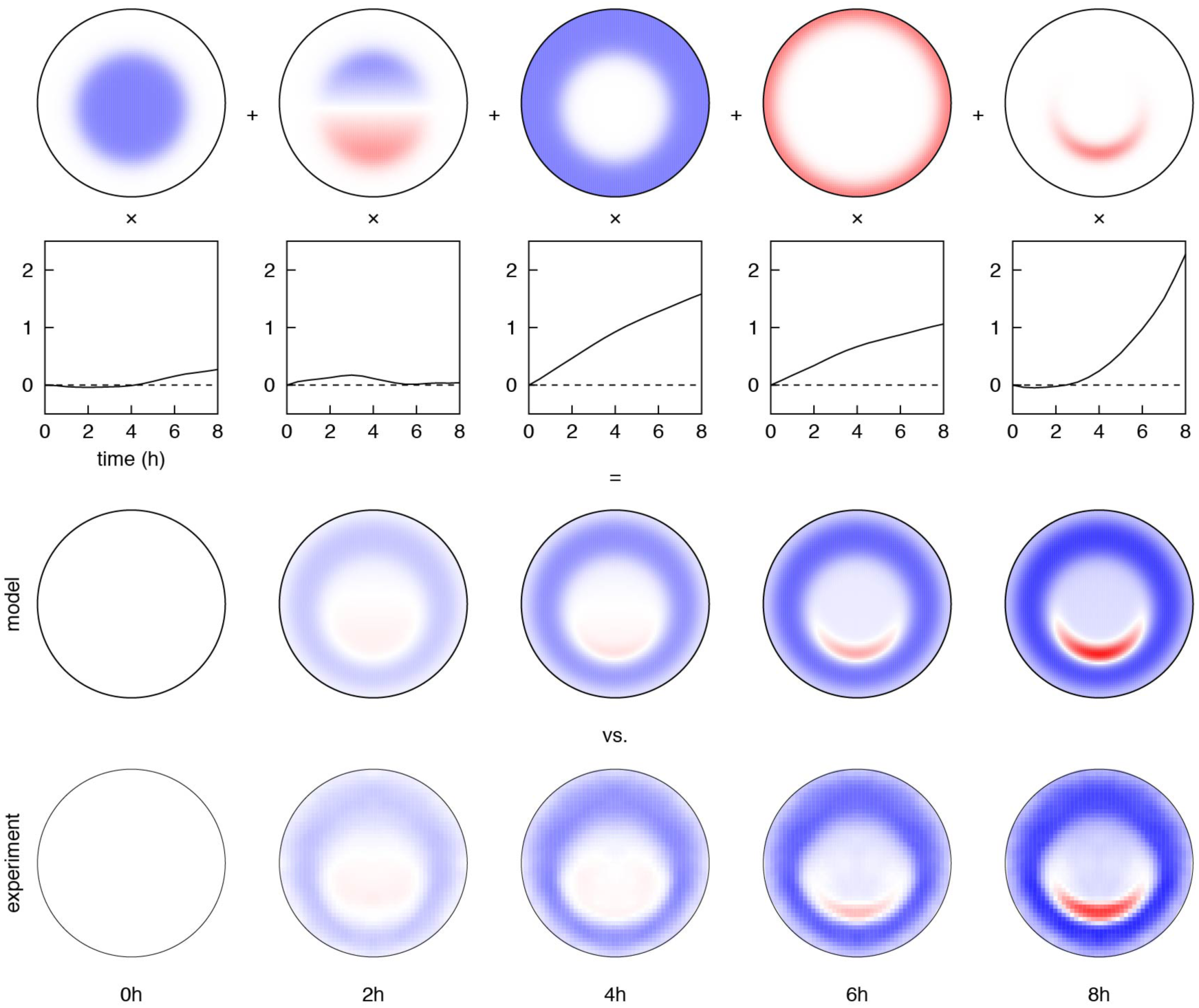
Detailed model for area changes. The log of area changes in the tissue is modeled as a linear combination of five modes, from left to right: uniform and graded expansion within the EP, EE expansion, reduced expansion near the tissue border, and areal contraction of the prospective primitive streak. Amplitudes vs. time are plotted below each mode, and the resulting area changes are compared to experimental area changes (from a symmetrized, average embryo) at successive times (colors as in Fig. 1c).

**Extended Data Figure 4.**
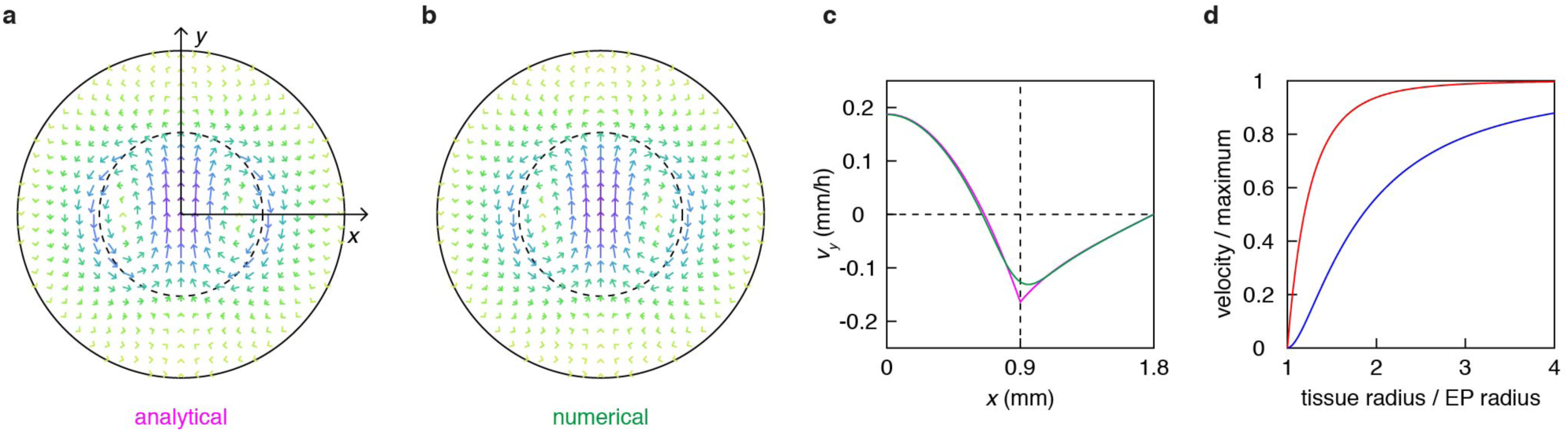
Analytical model. **a,** Velocity field computed analytically in the limit of a vanishing margin width, with a sinusoidal tension profile, *T* / *μ* = cos*θ*, cf. equation (S43) (dashed line, margin; solid line; tissue border). **b**, Velocity field computed numerically with the same tension profile but a finite margin width, *w* = 0.1 mm, cf. equation (S30). **c,** Comparison between the analytical and numerical velocity profiles as a function of distance to the center, showing that the two profiles differ appreciably only near the margin (dashed line); the slope becomes discontinuous in the limit of a vanishing margin width. **d**, Analytical curves showing how the global (“swimming”) velocity of the embryo (blue) and relative movement within the embryo (red) depend on the total size of the tissue relative to the EP. Motion within the embryo, thus the progress of gastrulation, is predicted to be much less sensitive to the presence of the tissue border, tending more rapidly to its maximum value.

**Extended Data Figure 5.**
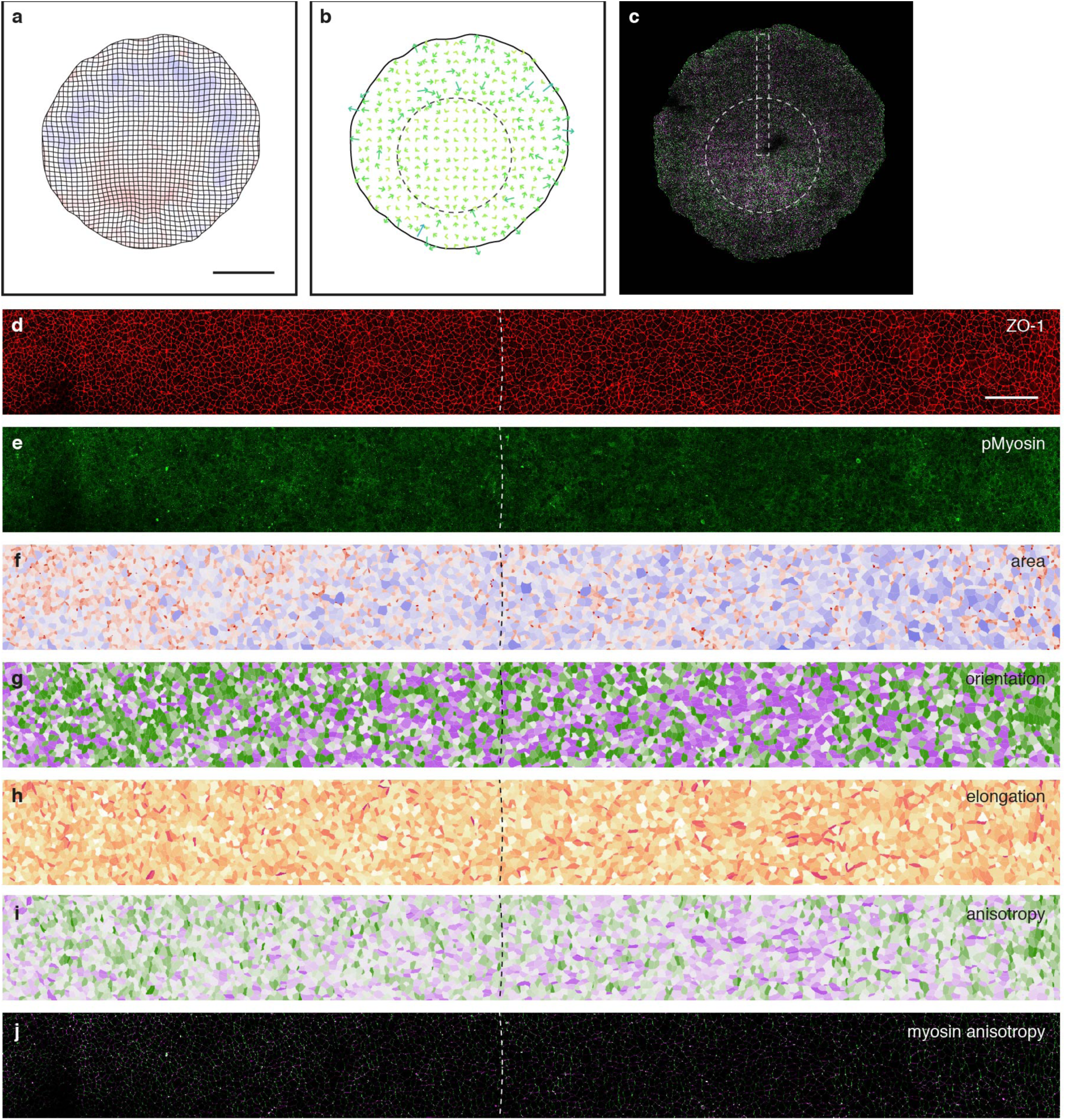
Quantification of cell shape and phosphorylated Myosin anisotropy in an anterior portion of an early embryo. **a**-**b,** Deformation map (**a**) and apparent forces (**b**; dotted line, margin) from live imaging of an early embryo (*t* ≈ *t*_0_), and junctional myosin in the same embryo after fixation (colors denote orientation; magenta, radial; green, orthoradial). **d**-**j,** ZO-1 (**d**) and phospho-Myosin (**e**) immunofluorescence used to segment and quantify cell areas (**f**), cell orientation (**g**), cell elongation (**h**), cell anisotropy (**i**) and junctional phosphoMyosin anisotropy (**j**; colors as in **c**) of the region boxed in **c** (see Supplementary Methods). Scale bars, **a**, 1 mm, **d**, 100 µm.

**Extended Data Figure 6.**
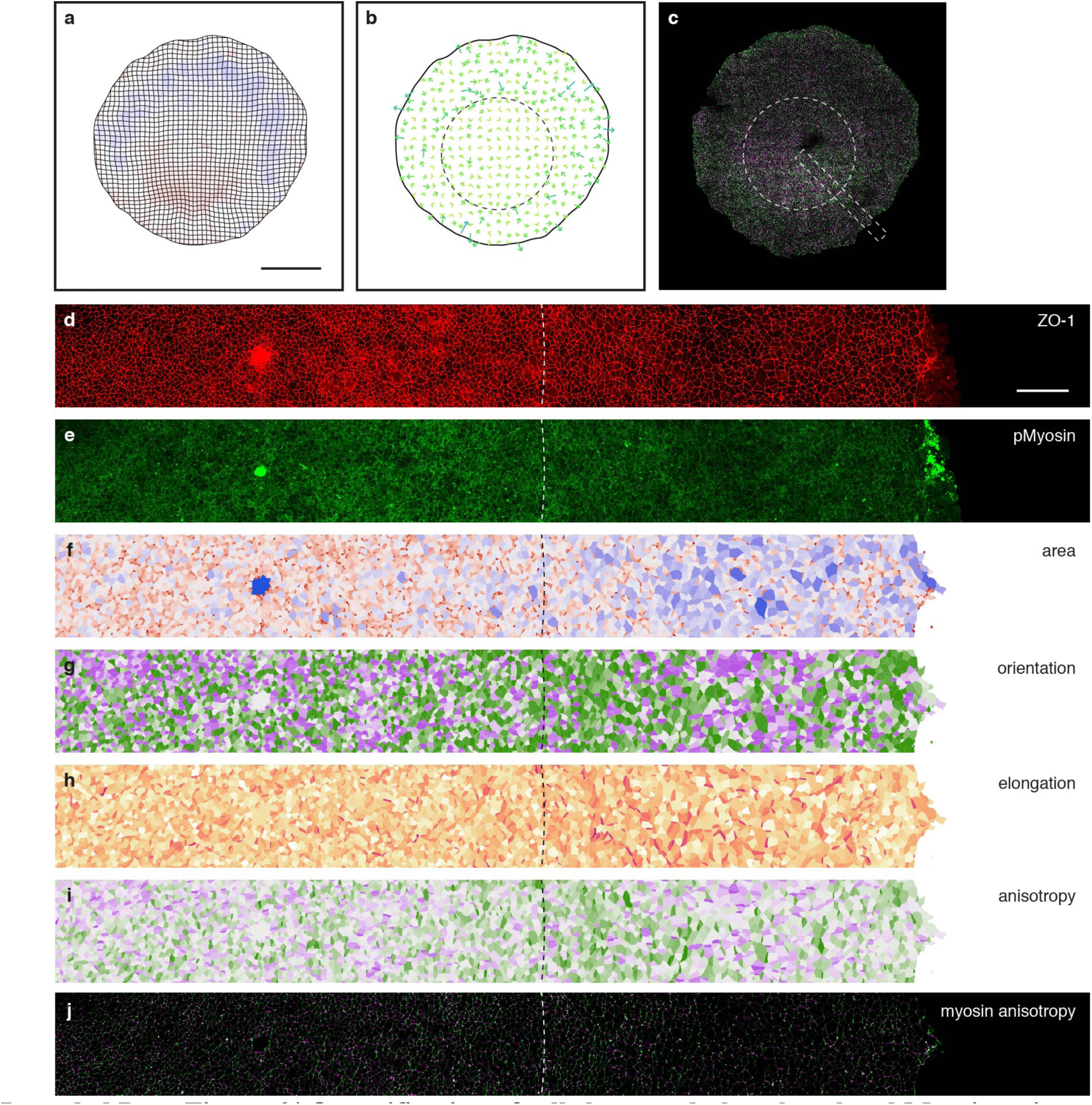
Quantification of cell shape and phosphorylated Myosin anisotropy in a posterior portion of an early embryo. **a**-**b,** Deformation map (**a**) and apparent forces (**b**; dotted line, margin) from live imaging of an early embryo (*t* ≈ *t*_0_), and junctional myosin in the same embryo after fixation (colors denote orientation; magenta, radial; green, orthoradial). **d**-**j,** ZO-1 (**d**) and phospho-Myosin (**e**) immunofluorescence used to segment and quantify cell areas (**f**), cell orientation (**g**), cell elongation (**h**), cell anisotropy (**i**) and junctional phosphoMyosin anisotropy (**j**; colors as in **c**) of the region boxed in **c** (see Supplementary Methods).

**Extended Data Figure 7.**
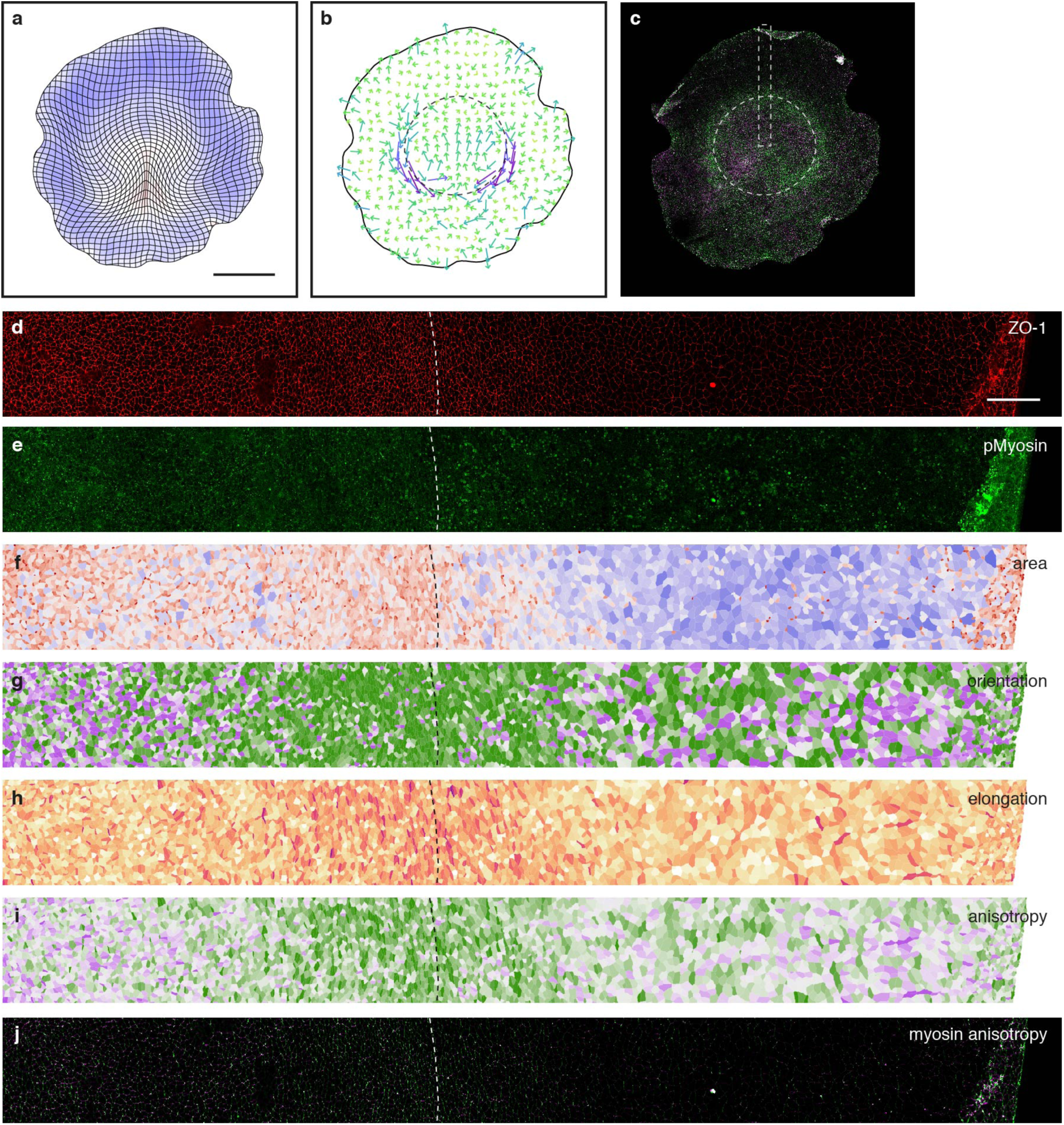
Quantification of cell shape and phosphorylated Myosin anisotropy in an anterior portion of a gastrulating embryo. **a**-**b,** Deformation map (**a**) and apparent forces (**b**; dotted line, margin) from live imaging of an early embryo (*t* ≈ *t*_0_ + 4 h), and junctional myosin in the same embryo after fixation (colors denote orientation; magenta, radial; green, orthoradial). **d**-**j,** ZO-1 (**d**) and phospho-Myosin (**e**) immunofluorescence used to segment and quantify cell areas (**f**), cell orientation (**g**), cell elongation (**h**), cell anisotropy (**i**) and junctional phosphoMyosin anisotropy (**j**; colors as in **c**) of the region boxed in **c** (see Supplementary Methods).

**Extended Data Figure 8.**
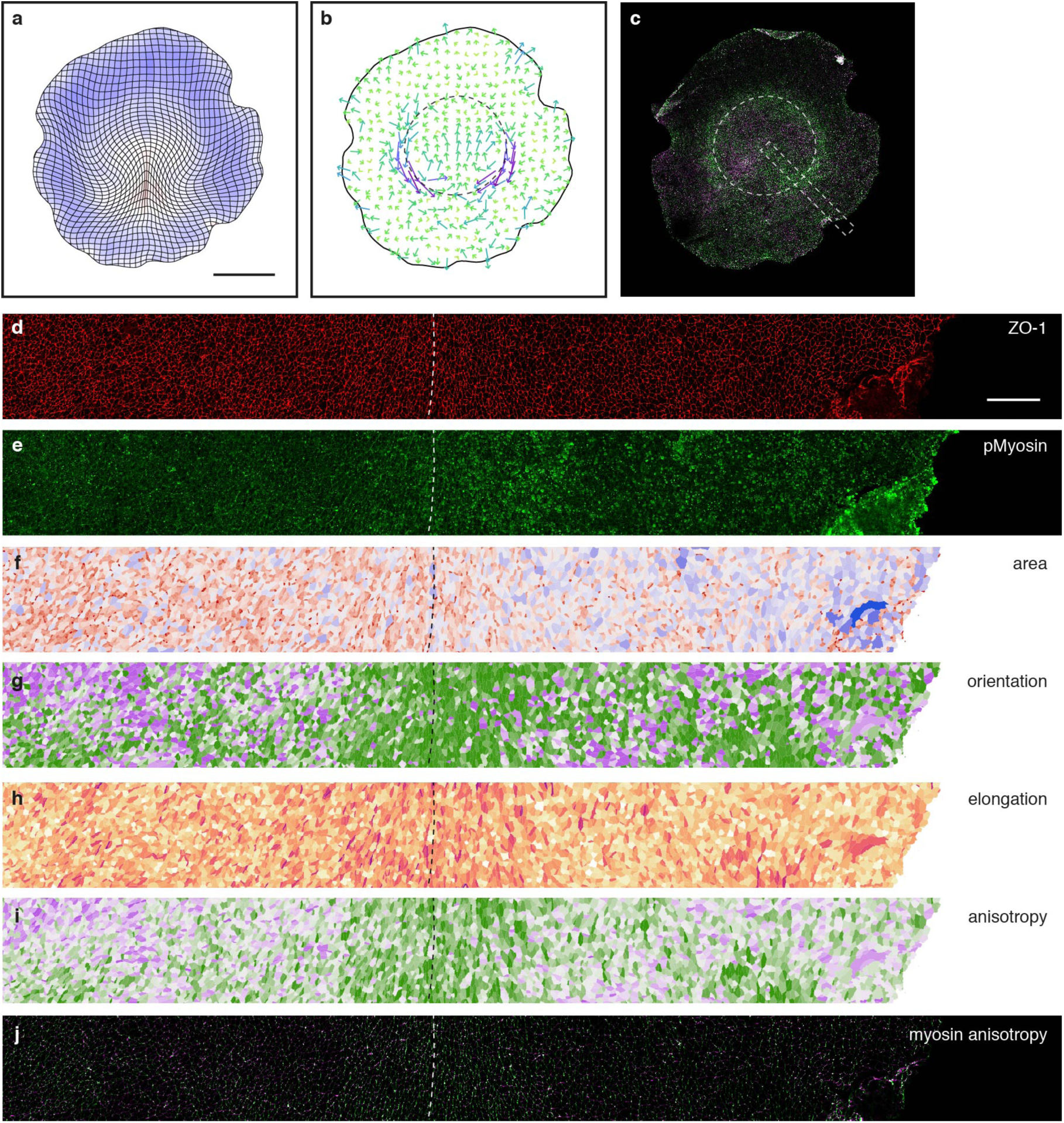
Quantification of cell shape and phosphorylated Myosin anisotropy in a posterior portion of a gastrulating embryo. **a**-**b,** Deformation map (**a**) and apparent forces (**b**; dotted line, margin) from live imaging of an early embryo (*t* ≈ *t*_0_ + 4 h), and junctional myosin in the same embryo after fixation (colors denote orientation; magenta, radial; green, orthoradial). **d**-**j,** ZO-1 (**d**) and phospho-Myosin (**e**) immunofluorescence used to segment and quantify cell areas (**f**), cell orientation (**g**), cell elongation (**h**), cell anisotropy (**i**) and junctional phosphoMyosin anisotropy (**j**; colors as in **c**) of the region boxed in **c** (see Supplementary Methods).

**Extended Data Figure 9.**
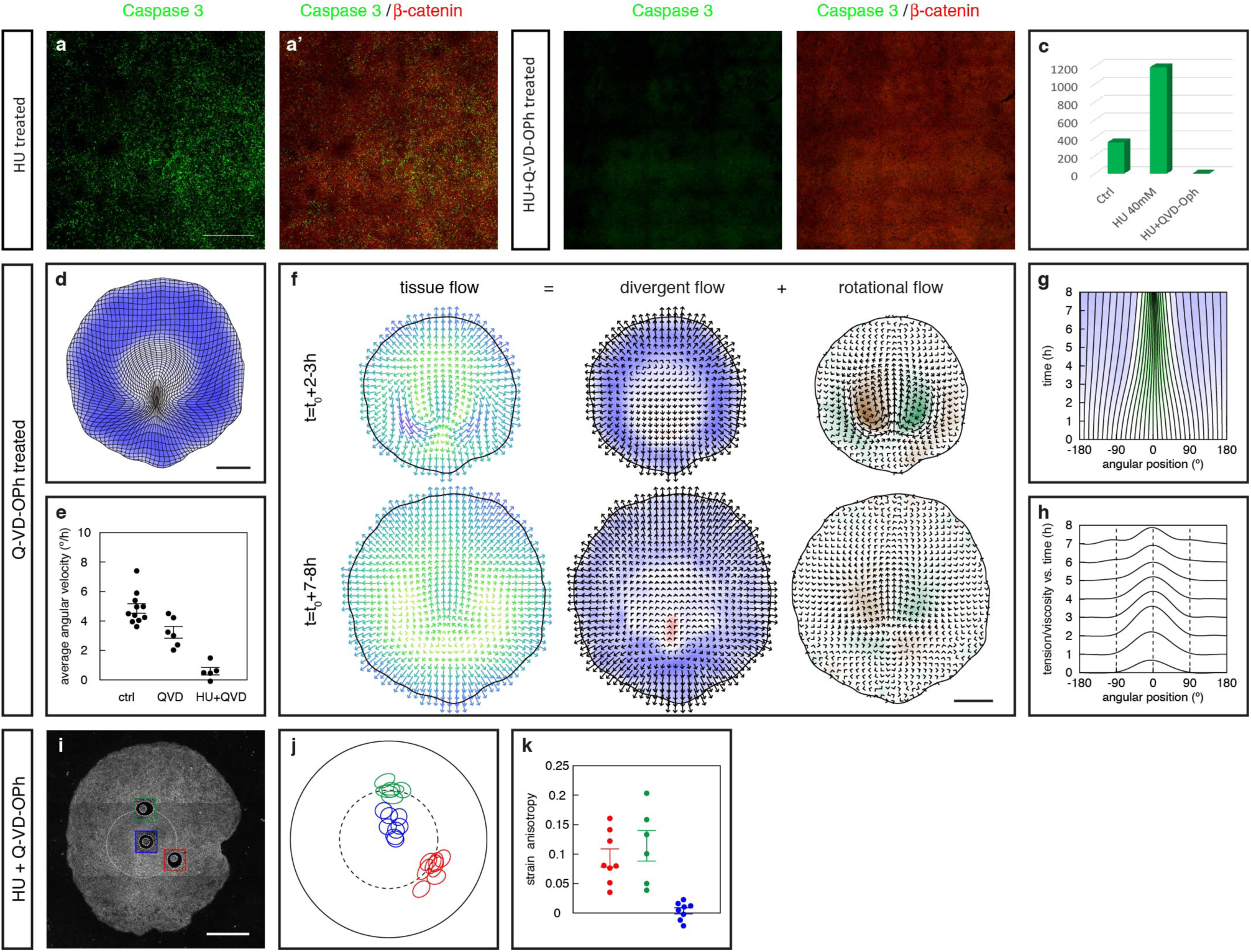
Characterization of the effect of cell division and apoptosis inhibition. **a**-**c**, Embryos treated with HU (**a**, **a’**) or with HU+Q-VD-OPh (**b**, **b’**), stained for caspase 3 (green), which labels apoptotic cells, and counterstained with beta-catenin (red). Quantification of the number of caspase 3 positive cells for each condition is shown in (**c**). **d**-**h**, Effect of Q-VD-OPh alone on gastrulation movements. **d**, Deformation map (average embryo; *n* = 5 embryos). **e**, Average angular velocity along the margin at *t* = *t*_0_ + 6.5-7.5 h. **f**, Decomposition of the tissue velocity field into divergent and rotational components (averages over embryos). **g**, Time evolution of angular positions (average embryo). **h**, Tension/viscosity profiles from model fit to average embryo. **i**-**k**, 250 μm circular laser cuts in a single memGFP embryo (**i**) and representation of all laser cut experiments (**j**) for which which the response was quantified (**k;** bars, mean ± SE) (red, posterior margin; green, anterior margin; blue, EP). Scale bars, **a**, 500 µm; **d**, **f**, **i**, 1 mm.

**Extended Data Table 1.**
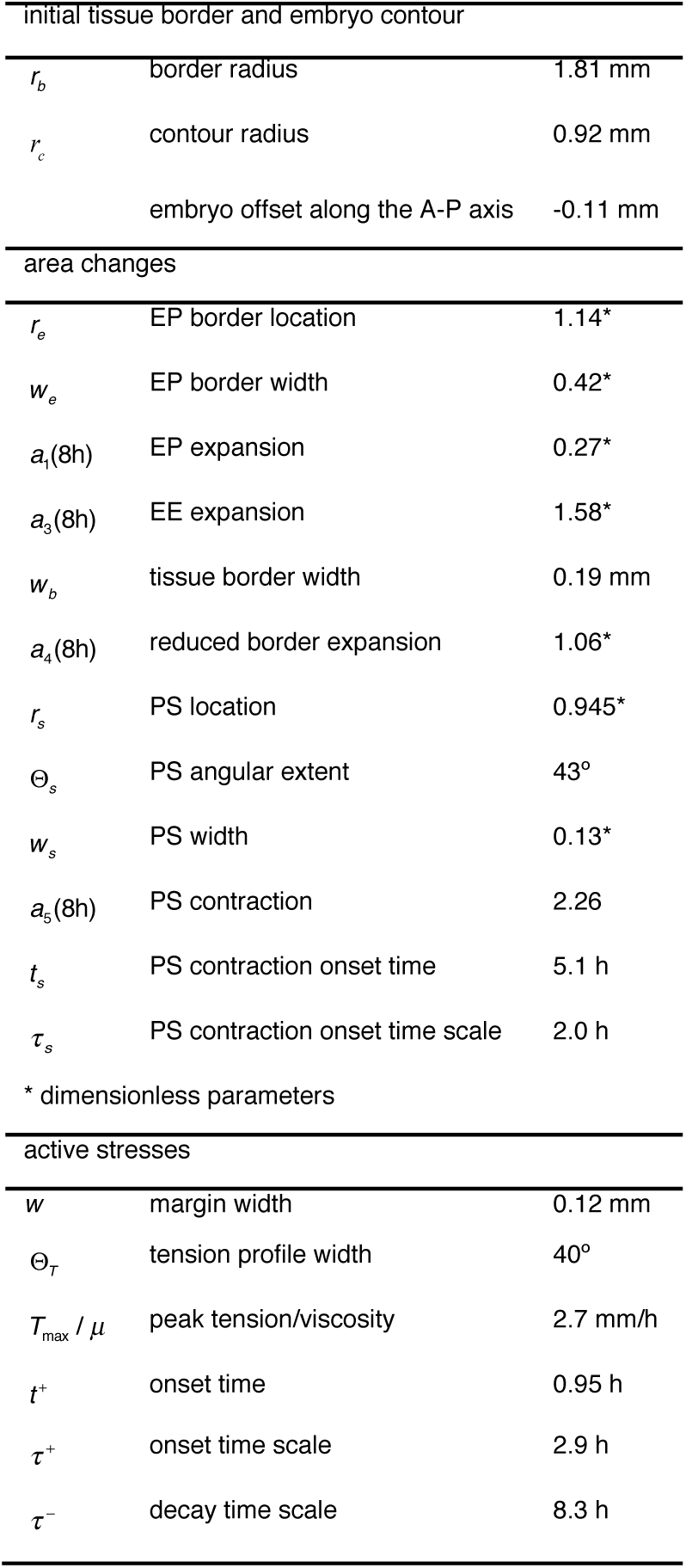
Model parameters for the synthetic embryo. The parameter values in this table were used to simulate the synthetic embryo of Fig. 2h, i and Supplementary Video 6, as described in Supplementary Methods.

| Initial tissue border and embryo contour |  |  |
| --- | --- | --- |
| $r_b$ | border radius | 1.81 mm |
| $r_c$ | contour radius | 0.92 mm |
|  | embryo offset along the A-P axis | -0.11 mm |
| area changes |  |  |
| $r_e$ | EP border location | 1.14* |
| $w_e$ | EP border width | 0.42* |
| $a_1(8h)$ | EP expansion | 0.27* |
| $a_3(8h)$ | EE expansion | 1.58* |
| $w_b$ | tissue border width | 0.19 mm |
| $a_4(8h)$ | reduced border expansion | 1.06* |
| $r_s$ | PS location | 0.945* |
| $\Theta_s$ | PS angular extent | 43° |
| $w_s$ | PS width | 0.13* |
| $a_5(8h)$ | PS contraction | 2.26 |
| $t_s$ | PS contraction onset time | 5.1 h |
| $\tau_s$ | PS contraction onset time scale | 2.0 h |
| * dimensionless parameters |  |  |
| active stresses |  |  |
| $w$ | margin width | 0.12 mm |
| $\Theta_T$ | tension profile width | 40° |
| $T_{\max} / \mu$ | peak tension/viscosity | 2.7 mm/h |
| $t^+$ | onset time | 0.95 h |
| $\tau^+$ | onset time scale | 2.9 h |
| $\tau^-$ | decay time scale | 8.3 h |

